# Development of attention-based robust deep learning model for tongue diagnosis by smartphone

**DOI:** 10.1101/2023.02.02.526804

**Authors:** Changzheng Ma, Chaofei Gao, Siyu Hou, Shao Li

## Abstract

Compared with tongue diagnosis using tongue image analyzers, tongue diagnosis by smartphone has great advantages in convenience and cost for universal health monitoring, but its accuracy is affected by the shooting conditions of smartphones. Developing deep learning models with high accuracy and robustness to changes in the shooting environment for tongue diagnosis by smartphone and determining the influence of environmental changes on accuracy are necessary. In our study, a dataset of 9003 images was constructed after image pre-processing and labeling. Next, we developed an attention-based deep learning model (Deep Tongue) for 8 subtasks of tongue diagnosis, including the spotted tongue, teeth-marked tongue, and fissure tongue et al, which the average AUC of was 0.90, higher than the baseline model (ResNet50) by 0.10. Finally, we analyzed the objective reasons, the brightness of the environment and the hue of images, affecting the accuracy of tongue diagnosis by smartphone through a consistency experiment of direct subject inspection and tongue image inspection. Finally, we determined the influence of environmental changes on accuracy to quantify the robustness of the Deep Tongue model through simulation experiments. Overall, the Deep Tongue model achieved a higher and more stable classification accuracy of seven tongue diagnosis tasks in the complex shooting environment of the smartphone, and the classification of tongue coating (yellow/white) was found to be sensitive to the hue of the images and therefore unreliable without stricter shooting conditions and color correction.

## Introduction

Tongue diagnosis was one of the most important diagnostic methods in TCM [1, 2]. Traditional Chinese Medicine (TCM) as alternative medicine was included in the Global Medical Compendium (2019 Edition) by the World Health Organization (WHO)[3]. In TCM, tongue diagnosis can visually demonstrate the health status.

Tongue diagnosis can provide an effective, non-invasive way to assist in health monitoring and the assessment of the patient’s physical condition. The irregular lifestyle of modern society can easily lead people to be in a sub-healthy state. Therefore, people’s demand for health monitoring is gradually increasing. Artificial intelligence techniques are widely used in biomedical fields, including research about tongue image analysis[4, 5]. The different characteristics of the tongue (such as color, shape, etc.) were significantly related to the health condition and different diseases [6-8], such as gastritis [9, 10], cancer[11–13], Corona Virus Disease-19 (COVID-19) [14, 15], diabetes[16–18]. Therefore, tongue diagnosis can be used for health monitoring and the prediction of diseases.

At present, tongue image analyzers (TIAI) are commonly used in tongue image photography but have some disadvantages that are difficult to overcome for health monitoring. For example, Huo et al. developed a tongue shape classification method with a data set of 318 tongue images by integrating image preprocessing and a convolution neural network (CNN) [19]. The accuracy rates of classification of fissure tongue, spotted tongue, and teeth-marked tongue were 98%, 90%, and 81.2% respectively. Hou et al. conducted research on the classification of tongue color based on CNN with a data set of 1500 tongue images, and the accuracy rate was 89% [20]. In addition, some scholars have conducted methodological studies for different subtasks of tongue diagnosis, such as tongue color, fissure, tongue shape, and tongue coating[21–27]. The above study achieved high accuracy of tongue diagnosis based on TIAI. However, TIAI as a tool for tongue image acquisition is large and costly, which leads to its applicability only to fixed places such as medical institutions and cannot meet the demand for people to monitor their health daily. Although it has the advantages of a good shooting effect and low interference, it is hard to popularize due to its high cost and inconvenience. Therefore, although TIAI could achieve intelligent tongue diagnosis with high accuracy, it is difficult to be applied to daily health monitoring.

Tongue diagnosis using smartphones can avoid the above problems and has enormous advantages. Smartphones are massively widespread in society and many studies have also focused on the great value of smartphones and personal smart devices for health monitoring [28–30]. The smartphone was more convenient and cheaper to take tongue images than TIAI, so it was suitable for public health monitoring and was of great significance to establishing health records and disease prevention.

However, the shooting environment of the smartphone is complex and open compared to TIAI, which leads to lower accuracy and higher indeterminacy. Therefore, it is necessary to develop classification models with high robustness for tongue images captured by smartphones and quantify the impact of environmental changes on the classification results to reduce indeterminacy.

Although there were some correction or quality assessment methods for tongue images to improve the classification accuracy of tongue diagnosis by smartphone, it is also necessary to analyze the influence of shooting conditions on the accuracy and develop more robust classification models. There are methods of correcting by comparing pictures before and after using flash to improve accuracy and methods of filtering out low-quality tongue images by deep learning [31, 32]. However, these methods cannot completely solve the influence of complex shooting conditions on tongue diagnosis. Therefore, it is necessary to systematically explore the main objective factors affecting the accuracy of tongue diagnosis and analyze their influence on the accuracy. The results can be used to assess the reliability of different tongue diagnosis subtasks, guide the optimization of shooting requirements, screen low-quality pictures and develop correction methods.

In this study, we constructed a tongue image dataset with a capacity of 9003, developed an attention-based classification model (Deep Tongue) for multi-task tongue diagnosis of tongue images taken by smartphone, and compared it with the baseline model. Then, a consistency comparison experiment between direct subject inspection and tongue image inspection was conducted to explore the main factors affecting tongue diagnosis by smartphone. Finally, we quantified the robustness of the Deep Tongue model to shooting conditions changes through simulation experiments in our dataset.

## Method

### Data collection and tongue image preprocessing

Tongue images were collected according to some rules by a smartphone APP. Subjects were provided with shooting instructions and shooting samples during image acquisition and asked to expose the complete flat tongue and to take the picture in a natural light environment or in a fluorescent environment to avoid the effects of brightness and color temperature. Photographs should be clear, with the tongue body occupying at least a quarter of the picture. The research plan for collecting tongue images was designed in accordance with the “Declaration of Helsinki” and was approved by the Human Ethics Committee of Institution Review Board of Tsinghua University(20200069). After explaining the nature of the study, we obtained verbal informed consent was obtained from participants. Then the low-quality images with blurry, distorted background color, or incomplete tongue were manually removed.

Tongues were segmented from complex backgrounds. TCM physicians labeled the complete tongue body of 180 tongue images with a square frame using “labelImg” software. We trained a tongue body recognition and locating model using the deep learning YOLOv5 model and 180 labeled tongue images [33]. Furthermore, using this model, we have carried out tongue body recognition and cutting on 9,003 tongue images, reshape to 224*224, and formed a pre-processed tongue body image data set. In this way, we can segment the tongue from the complex background to reduce the impact of the background on classification and improve accuracy.

Tongue images were labeled by TCM physicians. Two professional TCM physicians independently labeled spotted tongue, greasy coating, peeling fur, teeth-marked tongue, fissure tongue, tongue coating(yellow/white), tongue coating(thick/thin), and tongue color (crimson/red/pale/bluish purple) for tongue images. Brief introductions and explanations for the 8 tongue diagnosis subtasks were in Appendix 1. When the symptoms were not serious, the boundary between positive and negative samples was relatively vague, and there was inconsistency in labeling. Another TCM physician made a final judgment on images with inconsistent labels.

Eventually, the collected tongue images and labels together formed a dataset.

### Deep Tongue model development

In general, the abbreviated structure of the Deep Tongue model was as follows. First, the tongue image with the size of 224 * 224 was cut into 225 tiles, which were extracted features by the pre-trained ResNet50 model[34]. Then, the feature matrix was combined with the class embedding and added to the position embedding to obtain a new feature matrix. Secondly, the new feature matrix was transformed into an output feature matrix through the Deep Tongue block. Finally, the class embedding in the output feature matrix was as input to the final MLP classification module to obtain the final classification output (Figure 1).

**Figure 1.**
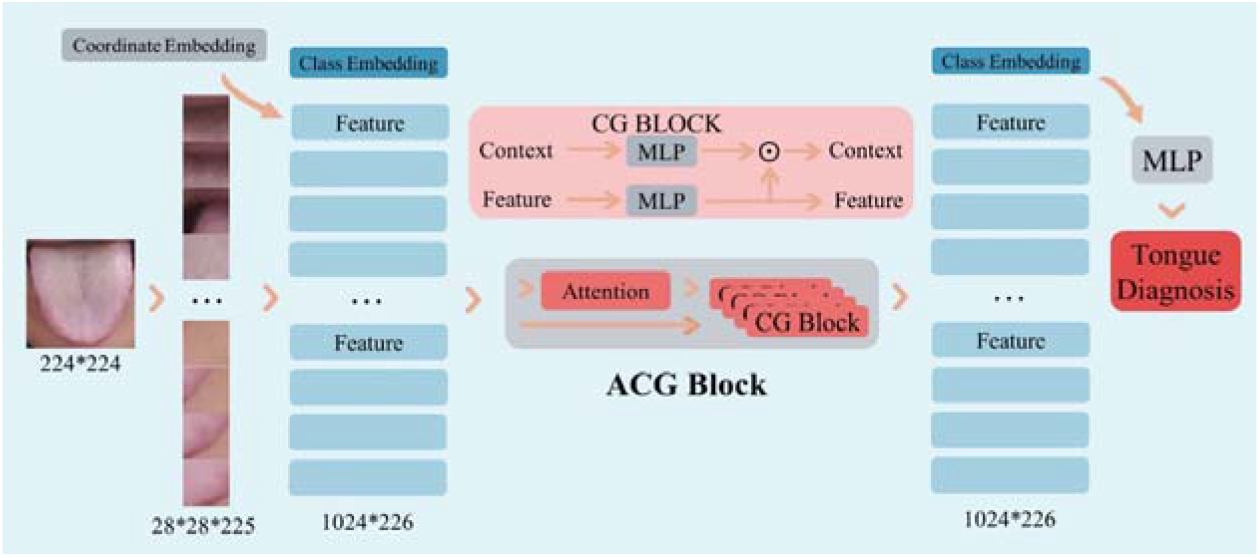
The structure flow chart of the Deep Tongue model.

#### Feature extraction module

The first part of the Deep Tongue model was the feature extraction module. The tongue image_∈(224 * 224)_ was cut into 225 tiles_∈(28*28)_ with a stride of 14. Each tile was converted into a feature vector_∈(1 * 2048)_ and extracted features by the pre-trained ResNet50 model. Then the 225 feature vectors were combined with a class embedding_∈(1 * 2048)_ to form a feature matrix_∈(226 * 2048)_. Finally, the feature matrix was added with the trainable coordinate embedding_∈(226*2048)_, which was only related to the position (abscissa and ordinate) of the tile on the tongue image. Coordinate embedding was a trainable feature vector that represents the coordinates of the location of the tile in the original image. This vector can introduce the relative and absolute position information of the features in the image into the model and improve the classification of the model.

In traditional convolutional neural network (CNN) methods, the convolutional process is the main part of training. In contrast, our approach used a CNN module, which has been pre-trained in many images, to extract features from the images. Further, we used these feature vectors for training and introduce an attention mechanism to enhance the model’s attention to key regions, thus improving the classification accuracy.

#### Attention-based module

The second part of the Deep Tongue model was an attention-based module. We input the preprocessed feature matrix S into an attention-based module to aggregate high-level representations from low-level representations [35]. As shown in formula (1), V_∈(1024 * 256)_, U_∈(1024 * 256)_ and W_∈(256 * 1024)_ was full connection layer. The output was a vector_∈(256 * 1)_, representing the weight of each feature. Output C was context, which represents the weighted sum of features. The sigmoid function was formula (2), which mapped inputs from 0 to 1 and normalized. The SoftMax function was formula (3). Like the sigmoid function, it mapped features to a value of 0 to 1, representing the weight value of each feature in the classification task. Features were introduced attention after this module, which was the key to achieving higher accuracy rates.

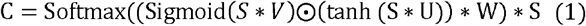

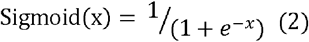

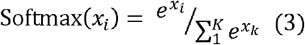

#### Context gating module

The third part of the Deep Tongue model was a context gating module. Context gating was adopted for introducing nonlinearity and strength discrimination into the input characteristics. As shown in formula (4), X was the input feature, and Y was the output feature. W was the trainable full connection layer and b was the bias. σ was the activation function.

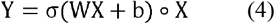

The context-gating calculation process included 4 cycles and each round was shown in the following formulas. The S matrix and the C vector were inputs and a new S matrix was the output. C_sum_ and S_sum_ were initialized as 0. MLP was a multilayer perceptron, which was a trainable module for the perceptron of features.

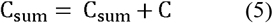

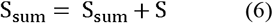

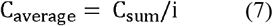

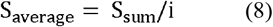

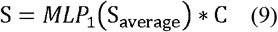

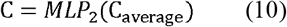

#### Classification module

The last part of the Deep Tongue model was a classification module. The class embedding of S was input to a trainable multilayer perceptron (MLP) layer module to obtain the final classification output. The final output vector was 1*2 or 1*4 for the 8 subtasks.

### Validation indicators and model comparison method

The training process and validation metrics were as follows. 5-fold cross-validation was adopted for training and the data set was divided into 5 parts on average, including 3 parts for training, 1 part for validation, and 1 part for testing. The data of different labels were imbalanced. Data of different labels were balanced into 1:1 in the training process to avoid large deviations. The learning rate was 10^−4^ and the loss function was cross-entropy. We adopted accuracy, sensitivity, specificity, F_β(β=2)_ score, the receiver operator characteristic (ROC) curve, and the area under the curve (AUC) as evaluation indicators.

The Deep Tongue model was compared with the baseline model of ResNet50 which was widely used as the baseline model for comparison in deep learning image classification problems and with good performance [36].

### Inter-rater consistency test for tongue diagnosis by smartphone and analysis reasons for the inconsistency

We calculated the inter-rater consistency of direct subject inspection and tongue image inspection by consistency test and analyzed the reasons for the inconsistency. TCM physicians did the direct subject inspection and tongue diagnosis and took tongue images for 551 people by smartphone. Then, TCM physicians made tongue image inspection judgments on the tongue images. After statistical analysis, we used consistency rate and Cohen’s Kappa to evaluate the consistency and analyzed the main objective reasons for the inconsistency by manual comparison.

### Robustness test for Deep Tongue model

We found that changes in the shooting environment would lead to a certain decrease in classification accuracy, which was manifested in the brightness and hue of tongue images. Therefore, we evaluated the robustness of the Deep Tongue model to their changes.

Hue and value were offset in HSV color space for tongue images to simulate hue and brightness change (Figure 2). The three dimensions of HSV color space were hue, saturation, and value. In the HSV color space, the hue range was H_□ [0,179]_. The simulation offset was ∇ H, and H=H±∇H, while H_□ [-4,4] *2_. Value was V_□ [0,255]_. The simulation offset was ∇V, and V=V±∇V, while ∇V_∈ [-4,4]*4_. The formulas for converting RGB color space to HSV was in Appendix 2.

**Figure 2.**
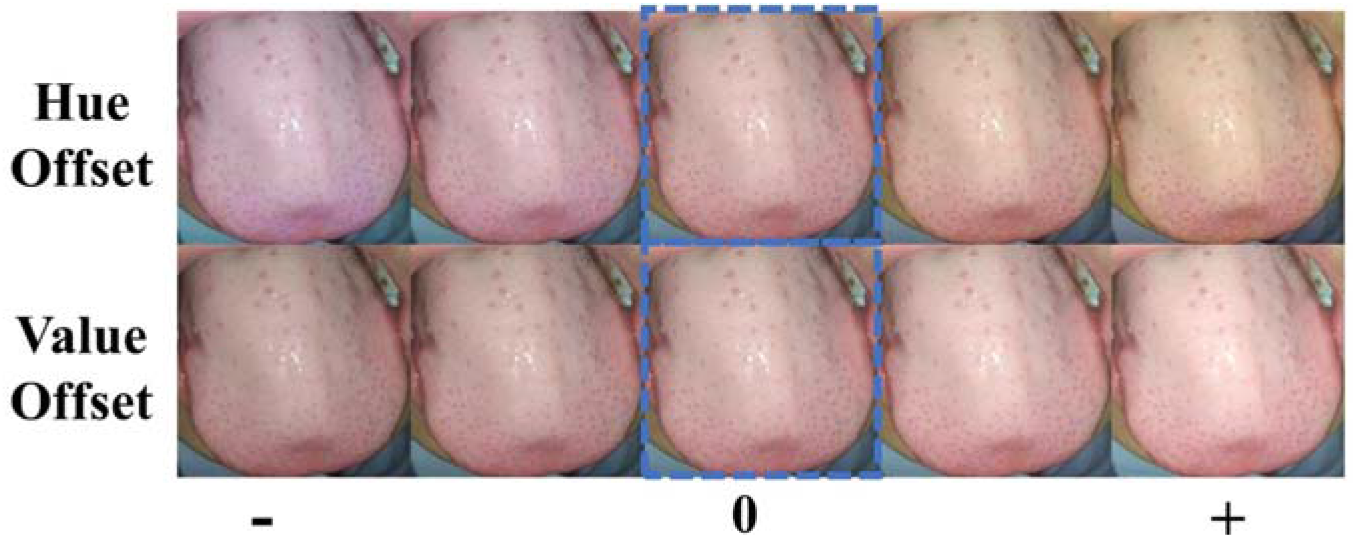
Examples of hue offset and value offset. The picture in the blue dashed box is the original.

## Results

First, we constructed a dataset with 9003 tongue images. Then, we proposed an attention-based classification model for 8 subtasks of tongue diagnosis and the average AUC was 0.90, which was significantly higher (P<0.01) than the ResNet50 model by 0.10. Finally, we analyzed the objective reasons affecting the accuracy of tongue diagnosis by smartphone through a consistency test of direct subject inspection and tongue image inspection and quantified the robustness of the Deep Tongue model to its changes through simulation experiments (Figure 3).

**Figure 3.**
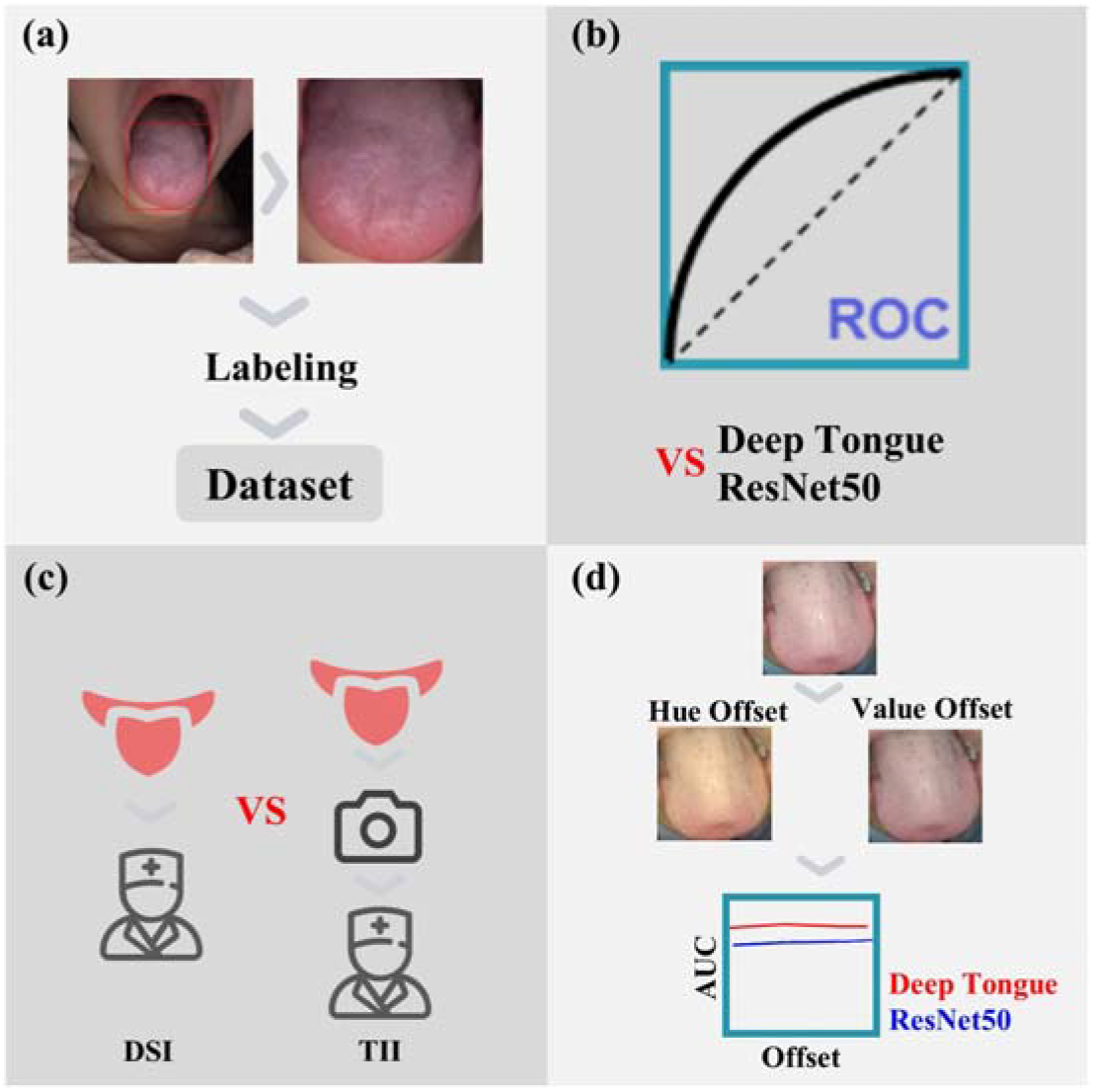
The overall design of our study. (a) dataset development (b) validation and model comparison (c) consistency test (d) robustness evaluation DSI: direct subject inspection, TII: tongue image inspection.

### Dataset development

We used smartphones to collect tongue images and filter out low-quality images (Figure 4). The tongue images were labeled with tongue diagnosis tags by professional TCM practitioners and the tongue body parts were identified and cut out from the tongue images by a trained YOLOv5 deep learning model to form a tongue dataset with a quantity of 9003 (Figure 5).

**Figure 4.**
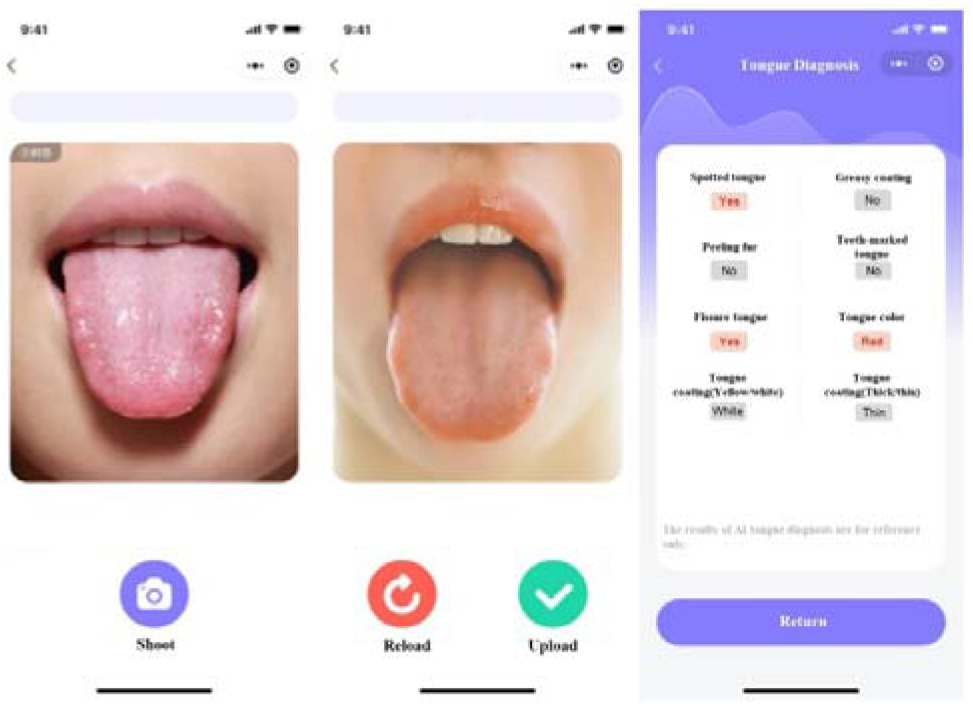
The operation interface of the smartphone app for tongue diagnosis.

**Figure 5.**
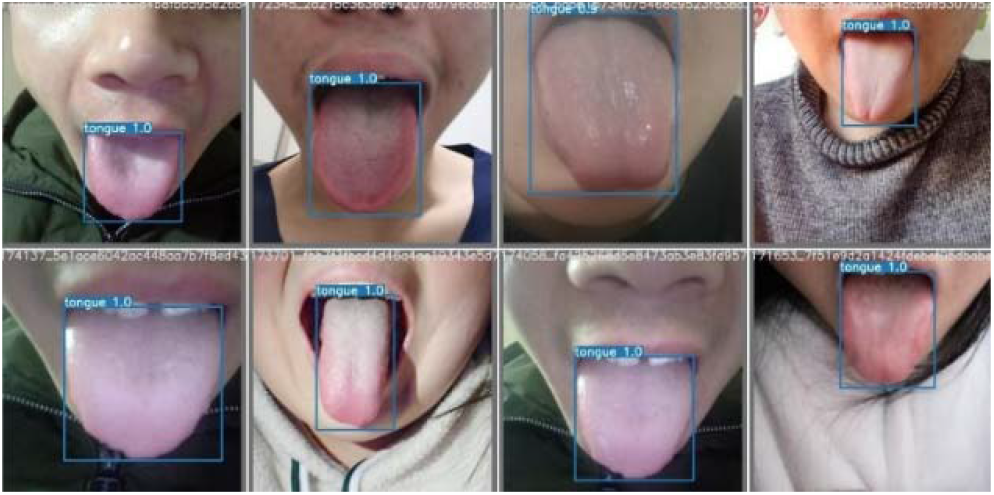
The samples of tongue body recognition and locating results.

Introductions and quantities for the 8 tongue diagnosis subtasks. Spotted tongue: red spots on the tongue surface, 73:8930; Greasy coating: greasy mucus adheres to the surface of the tongue, 985:8018; Peeling fur: incomplete peeling of tongue coating, 27:8976; Teeth-marked tongue: teeth indentation on the edge of the tongue, 322:8681; Fissure tongue: cracks on tongue body, 1782:7221; Tongue coating (yellow/white): yellow/white tongue coating on the tongue surface, 106:8897; Tongue coating (thick/thin): thick/thin tongue coating on the tongue surface, 1496:7507; Tongue color (crimson/red/pale/bluish purple): the color of tongue body,309:8399:290:5 (Figure 6).

**Figure 6.**
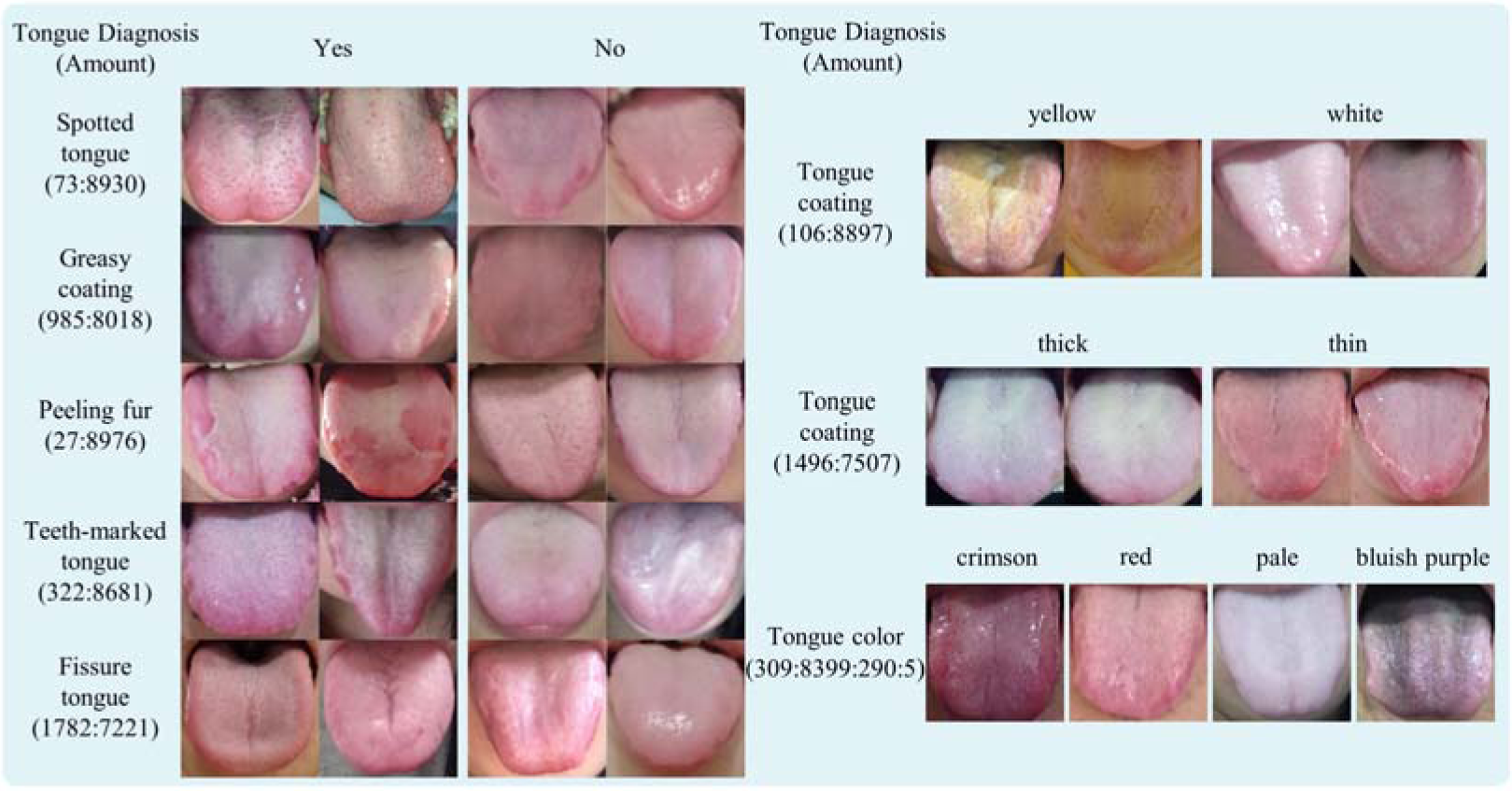
Examples and quantity of different tongue diagnosis subtasks.

### Validation and model comparison

The average AUC of the Deep Tongue model was 0.90, significantly higher than that of the ResNet50 model by 0.10. For the 8 subtasks, the AUCs of the two models were: spotted tongue: 0.83,0.63; greasy coating: 0.93,0.86; peeling fur:0.81,0.61; teeth-marked tongue: 0.94,0.73; fissure tongue: 0.95,0.94; tongue coating (yellow/white): 0.91,0.89; tongue coating (thick/thin): 0.90,0.83; tongue color (crimson/red/pale/bluish purple): 0.92,0.91. For each subtask, the AUC of Deep Tongue was higher than that of the ResNet50 model (Figure 7). In addition, the accuracy and F_β(β=2)_scores of the Deep Tongue model were higher than that of the ResNet50 model in all 8 subtasks.

**Figure 7.**
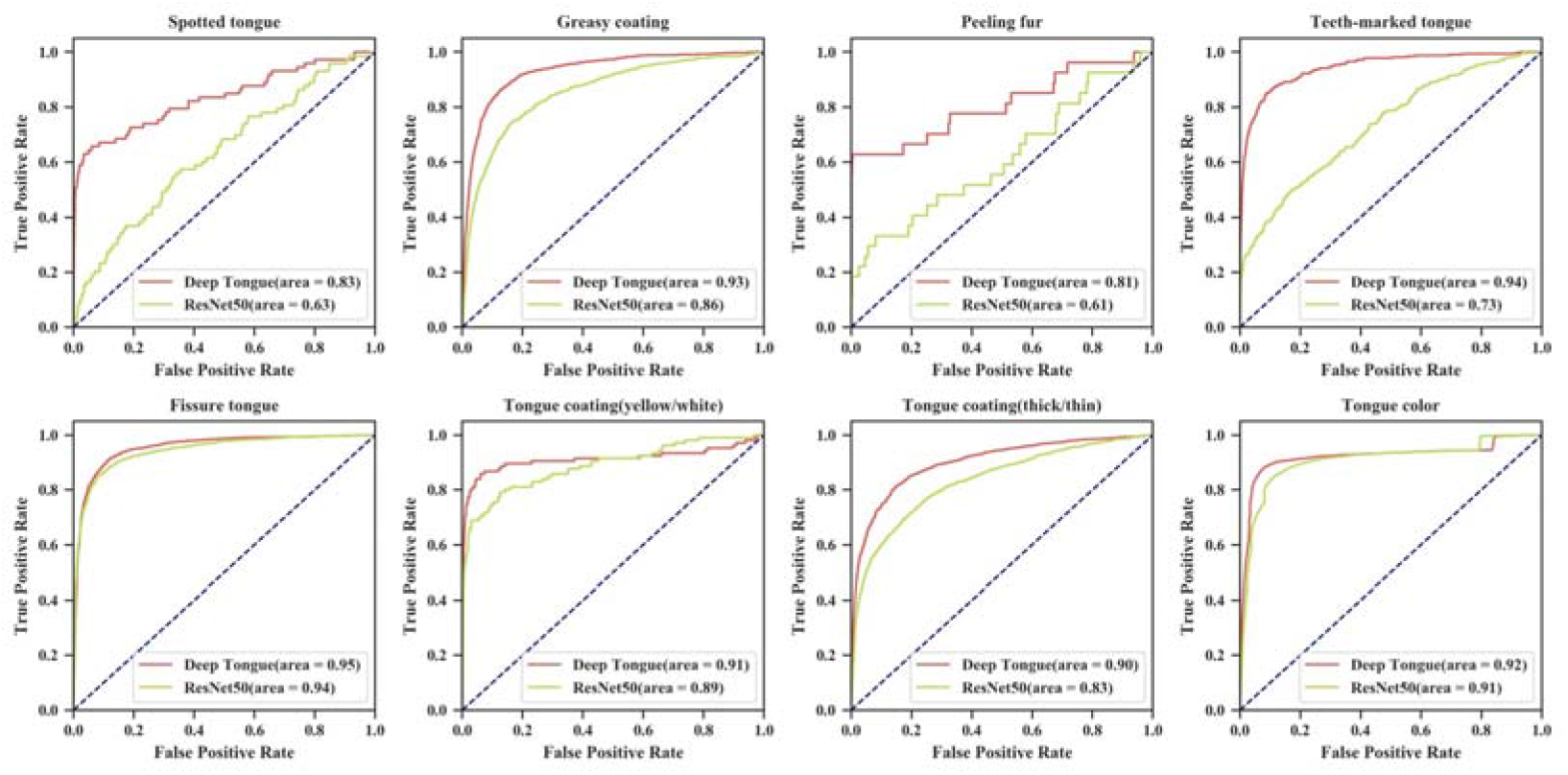
ROC curve and AUC for 8 tasks of Deep Tongue and ResNet50 models.

We compared the accuracy, sensitivity, specificity, and F_β(β=2)_score of the Deep Tongue model and baseline model. The thresholds of classification models were selected at the highest point of F_β(β=2)_ scores (Table 1).

**Table 1.** The accuracy, sensitivity, specificity, and F_β(β=2)_score of Deep Tongue and ResNet50 models for 8 subtasks.

| Method | Deep Tongue |  |  |  | ResNet50 |  |  |  |
| --- | --- | --- | --- | --- | --- | --- | --- | --- |
| | Acc | Spe | Sen | $F_{\beta}(\beta=2)$ | Acc | Spe | Sen | $F_{\beta}(\beta=2)$ |
| Spotted tongue | 0.99 | 0.99 | 0.49 | 0.48 | 0.97 | 0.97 | 0.11 | 0.05 |
| Greasy coating | 0.89 | 0.90 | 0.83 | 0.74 | 0.83 | 0.85 | 0.73 | 0.61 |
| Peeling fur | 1.00 | 1.00 | 0.52 | 0.60 | 0.99 | 1.00 | 0.15 | 0.17 |
| Teeth-marked tongue | 0.96 | 0.97 | 0.72 | 0.65 | 0.89 | 0.91 | 0.38 | 0.29 |
| Fissure tongue | 0.89 | 0.88 | 0.91 | 0.85 | 0.89 | 0.90 | 0.87 | 0.82 |
| Tongue coating (yellow/white) | 0.99 | 0.99 | 0.74 | 0.64 | 0.99 | 0.99 | 0.51 | 0.51 |
| Tongue coating (thick/thin) | 0.85 | 0.85 | 0.81 | 0.73 | 0.75 | 0.75 | 0.77 | 0.64 |
| Tongue color | 0.90 | 0.94 | 0.71 | 0.69 | 0.89 | 0.92 | 0.61 | 0.61 |

### Consistency of direct subject inspection and tongue image inspection and analysis of factors of inconsistency

The consistency between direct subject inspection and tongue image inspection was good for the average consistency of the 8 subtasks was 95.2% and the average Cohen’s Kappa was 0.83 (Table 2). The numbers of different labels for each subtask were shown in Figure 8.

**Table 2.**
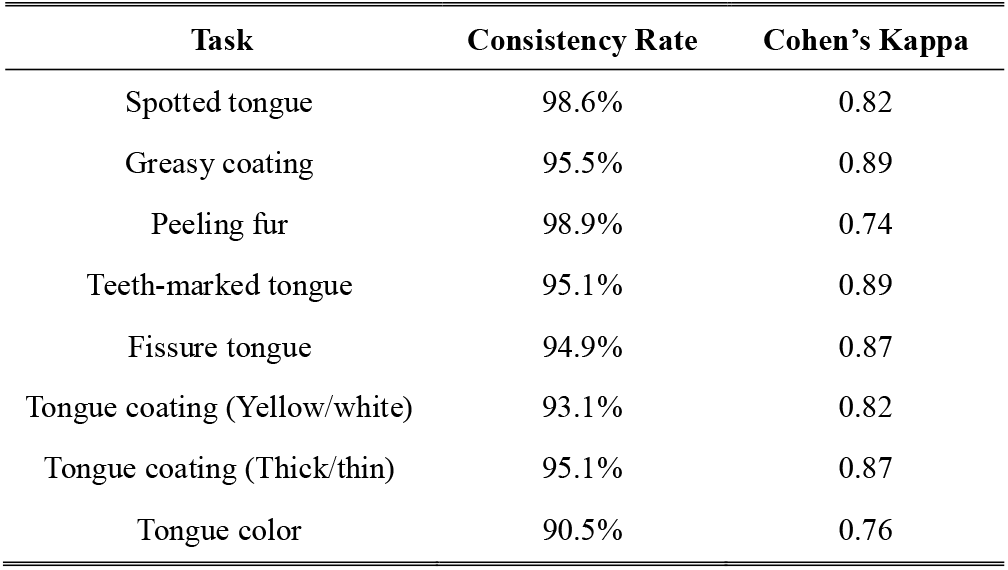
The consistency test result of consistency rate and Cohen’s Kappa.

| Task | Consistency Rate | Cohen's Kappa |
| --- | --- | --- |
| Spotted tongue | 98.6% | 0.82 |
| Greasy coating | 95.5% | 0.89 |
| Peeling fur | 98.9% | 0.74 |
| Teeth-marked tongue | 95.1% | 0.89 |
| Fissure tongue | 94.9% | 0.87 |
| Tongue coating (Yellow/white) | 93.1% | 0.82 |
| Tongue coating (Thick/thin) | 95.1% | 0.87 |
| Tongue color | 90.5% | 0.76 |

**Figure 8.**
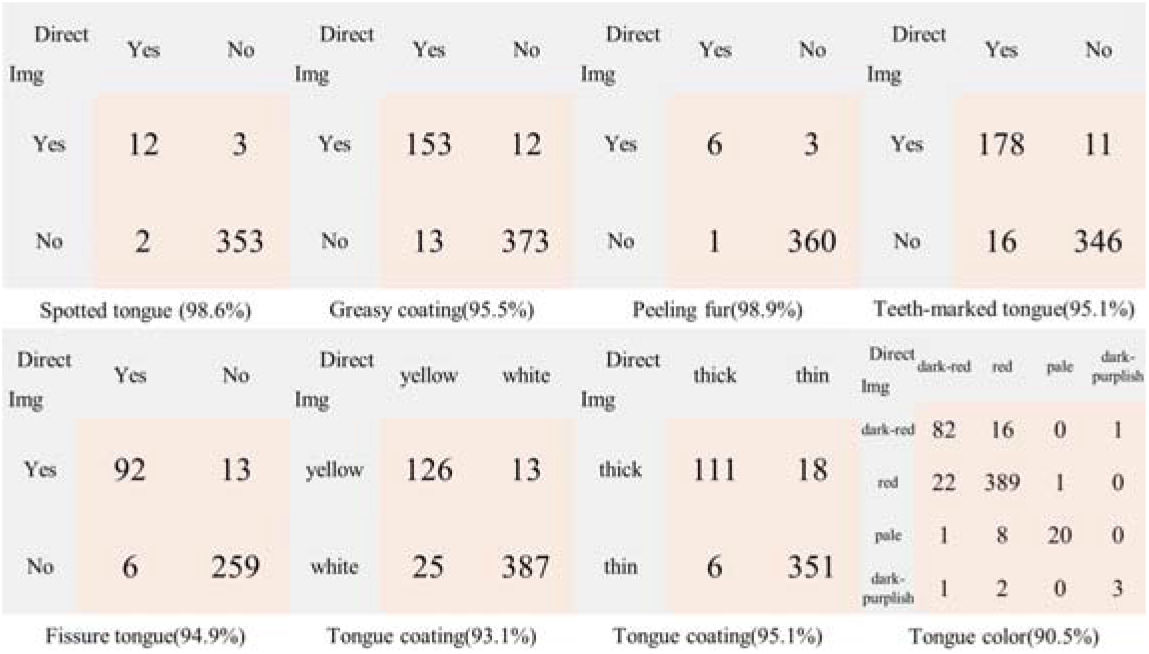
The consistency test results and accuracy rate of direct subject inspection and tongue image inspection tongue diagnosis. The abscissa was the labeling result of “direct subject inspection”, while the ordinate was the labeling result of “tongue image inspection”. The number part in the figure indicated the number of the label results.

Through manual analysis of the inconsistent data, we found that in addition to the subjective reasons of TCM physicians for tongue diagnosis, the main factors causing the inconsistency were the brightness of the environment and the hue of images. After the consistency analysis, based on the results of the direct subject inspection, the TCM physicians analyzed the main causes of the inconsistency between the tongue image inspection and the results of the direct subject inspection. The results showed that the brightness of the environment accounted for 22% of the total inconsistency, with 60% of the spotted tongue and 37% of the tongue-marked tongue. About 35% of the differences were caused by the hue of the images, among which 66% were caused by tongue coating (yellow/white) and 62% by tongue color (Table 3).

**Table 3.** Quantity and proportion of the main environmental factors causing the inconsistency: the brightness of the environment and the hue of the images.

| Task | Inconsistent | Brightness | Hue |
| --- | --- | --- | --- |
| Spotted tongue | 5 | 3(60%) | 0(0%) |
| Greasy coating | 25 | 8(32%) | 1(4%) |
| Peeling fur | 4 | 0(0%) | 0(0%) |
| Teeth-marked tongue | 27 | 10(37%) | 2(7%) |
| Fissure tongue | 19 | 2(11%) | 1(5%) |
| Tongue coating (Yellow/white) | 38 | 6(16%) | 25(66%) |
| Tongue coating (Thick/thin) | 24 | 8(33%) | 7(29%) |
| Tongue color | 52 | 5(10%) | 32(62%) |

### Robustness of Deep Tongue model

The results of the consistency test showed that the main factors affecting the accuracy of tongue diagnosis by smartphone were the brightness of the environment and the hue of images. We tested the robustness of the model by simulating the changes of the two main factors by changing the value and hue of images and the Deep Tongue model was more robust to the changes of value and hue than the ResNet50 model.

The AUCs of 7 subtasks remained stable with a slight reduction when the hue shifted while the AUC of tongue coating color decreased significantly when the hue was negatively shifted (Figure 9). The main reason for the obvious decrease was that the recognition of tongue coating color was greatly affected by the change of hue compared with other classification subtasks. Overall, the Deep Tongue model was less robust in the tongue coating(yellow/white) subtask and more robust in the other subtasks against the changes in the hue.

**Figure 9.**
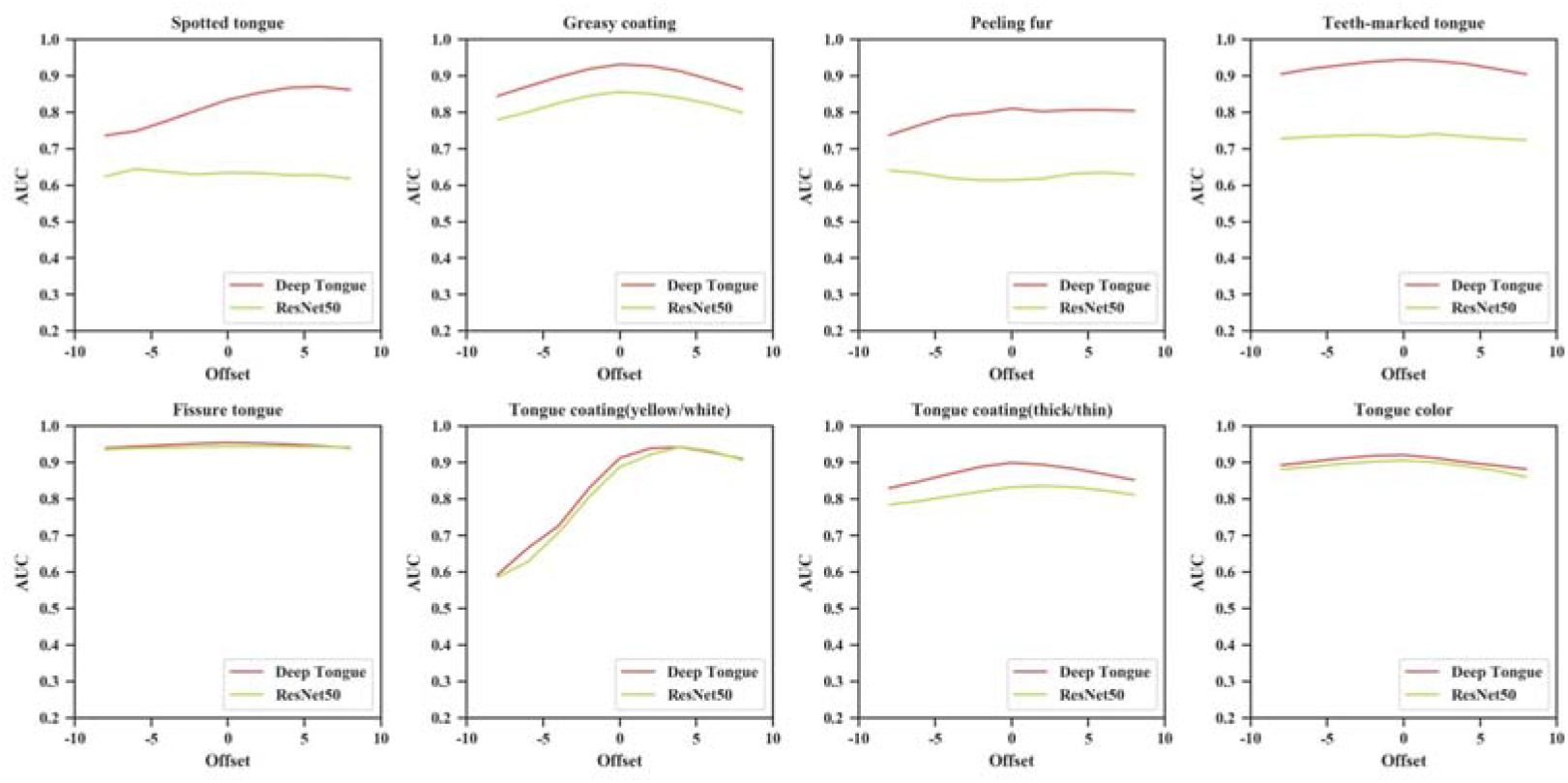
The robustness test result of ROC curves and AUCs for hue offset.

The AUCs of all 8 subtasks were greatly stable when the value shifted and the Deep Tongue model was more robust than the ResNet50 model in all subtasks against the changes in the value (Figure 10).

**Figure 10.**
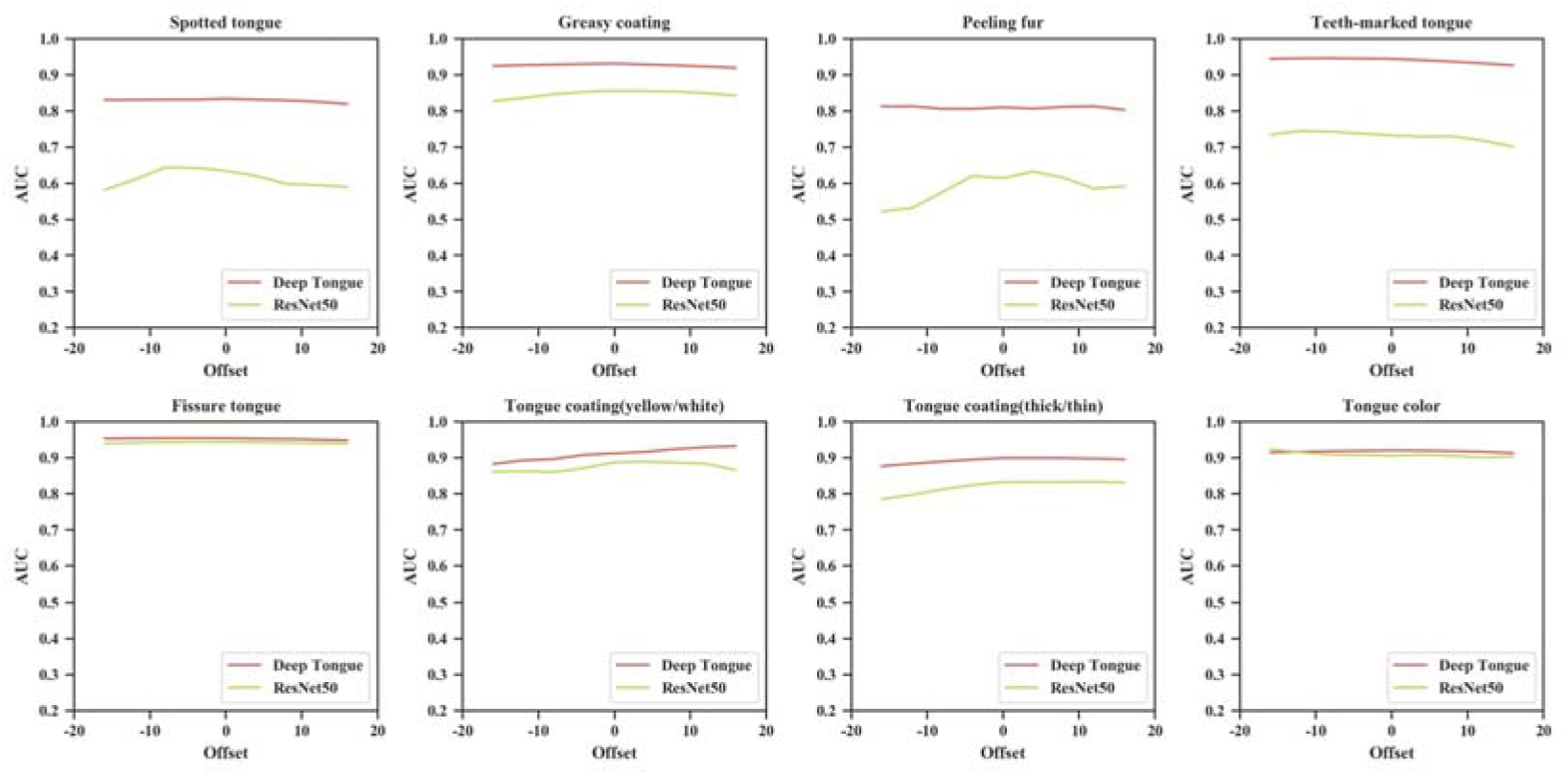
The robustness test result of ROC curves and AUCs for value offset.

Overall, we found that the deep Tongue model achieved higher accuracy and robustness to hue and value changes than the baseline model. At the same time, we also found that a small range of brightness changes had little impact on classification, the hue had little impact on seven tongue diagnosis tasks, but the tongue coating (yellow/white) was sensitive to hue changes. Therefore, we believed that Deep Tongue could achieve the reliable classification of tongue diagnosis in the complex shooting environment of the smartphone but the reliable classification of tongue coating (yellow/white) needs to be carried out under stricter shooting conditions and color correction.

## Discussion

Aiming at the need for smartphone-assisted tongue diagnosis in the open background, the attention-based module was introduced to construct a multi-task tongue diagnosis classification model for tongue images taken by smartphones and we analyzed the objective reasons affecting the accuracy of tongue diagnosis by smartphone through a consistency experiment of direct subject inspection and tongue image inspection and quantified the robustness of the Deep Tongue model to its changes through simulation experiments. The Deep Tongue model achieved higher and stable classification accuracy of seven tongue diagnosis tasks under the circumstance of small deviation of brightness and hue and the classification of tongue coating (yellow/white) was found to be sensitive to the hue of the images and therefore unreliable if the hue was not correctly corrected. In addition, the results of the robustness analysis can also be used to guide the optimization of shooting requirements, screen low-quality images, and develop correction methods. In summary, the work was a deeper exploration of automatic tongue diagnosis and an important step in promoting smartphone-assisted tongue diagnosis for health monitoring.

The Deep Tongue model achieved higher accuracy mainly for the attention mechanism was introduced to make the model has a higher perception of local key features. By introducing location information, the model effectively captured high-scale spatial features when focusing on low-scale features.

The significance of consistency testing is not only to analyze the objective causes of inconsistency but also to assess the degree of reliability of tongue diagnosis using images. TCM tongue diagnosis is a subjective evaluation index, and different doctors may give different diagnoses for the same patient. There is a loss of information and discrepancy in the photographed images compared to face-to-face tongue diagnosis, and this loss and discrepancy are unavoidable even with the use of a tongue scanner. The basis trying to avoid such discrepancies when taking the images, we also need to make an objective evaluation of the consistency to assess the degree of reliability of tongue diagnosis using images. Our results show that the consistency is high relative to the accuracy of current tongue diagnosis classification models.

We developed an APP to provide direct access to health monitoring (Figure 2). The use of an APP as a carrier greatly improves the efficiency of screening. We applied intelligent tongue diagnosis technology to the study of stomach disease screening. The oral cavity and the stomach are connected and belong to the same digestive system. In TCM, tongue diagnosis is an important tool for diagnosing gastric diseases. We tried to introduce the intelligent tongue diagnosis system into the monitoring and screening of gastric diseases and achieved good results. In that study, we found that tongue coating, fissure, and greasy were risk factors for precancerous lesions of the stomach. The introduction of these risk indicators can effectively improve the screening efficiency of gastric precancerous lesions.

Of course, our study has limitations. In the robustness analysis, we found that the tongue coating (yellow/white) was greatly affected by hue, which would lead to low reliability of its classification in practical applications. In the next work, we will focus on solving the problem of color correction in complex lighting environments.

## Appendix 1. Brief introductions and explanations for the 8 tongue diagnosis subtasks.

Spotted tongue: Red spots on the tongue surface. Heat syndrome. Greasy coating: Greasy mucus adheres to the surface of the tongue. Insufficiency-cold. Peeling fur: Incomplete peeling of tongue coating. Insufficiency of the spleen. Teeth-marked tongue: Teeth indentation on the edge of tongue. Deficiency of vital energy. Fissure tongue: Cracks on tongue body. Deficiency of essence and blood. Tongue coating (yellow/white): Yellow/white tongue coating on the tongue surface. Yellow: heat syndrome. Tongue coating (thick/thin): Thick/thin tongue coating on the tongue surface. Tongue color (crimson/red/pale/bluish purple): The color of tongue body. Excessive heat/ Heat syndrome/Insufficiency-cold/Blood stasis syndrome.

## Appendix 2. The formulas for converting RGB color space to HSV.

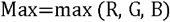

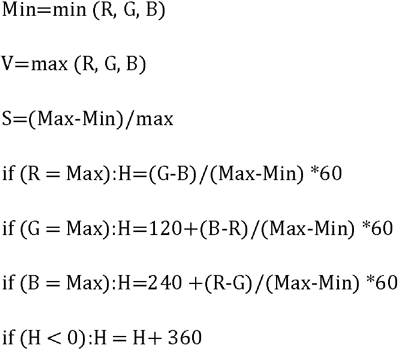

## Acknowledgment

Thanks to the doctors at Yijishan Hospital of Wannan Medical College for collecting tongue images and labeling tongue images.

## Reference

1. Li S: Mapping ancient remedies: Applying a network approach to traditional Chinese medicine. Science 2015, 350:S72–S74.

2. Su SB, Lu A, Li S, Jia W: Evidence-Based ZHENG: A Traditional Chinese Medicine Syndrome. Evid Based Complement Alternat Med 2012, 2012:246538.

3. Cyranoski D: Why Chinese medicine is heading for clinics around the world. Nature 2018, 561(7724):448–450.

4. Lei Y, Li S, Liu Z, Wan F, Tian T, Li S, Zhao D, Zeng J: A deep-learning framework for multi-level peptide-protein interaction prediction. Nat Commun 2021, 12(1):5465.

5. Zhou W, Yang K, Zeng J, Lai X, Wang X, Ji C, Li Y, Zhang P, Li S: FordNet: Recommending traditional Chinese medicine formula via deep neural network integrating phenotype and molecule. Pharmacol Res 2021, 173:105752.

6. Wang SJ, Xu ZX, Wang YQ: Tongue Characteristics of Different TCM Syndromes on Asthma. Adv Bio Sci Res 2016, 3:202–205.

7. Ye J, Cai X, Cao P: Problems and prospects of current studies on the microecology of tongue coating. Chin Med 2014, 9(1):9.

8. Chen H, Li Q, Li M, Liu S, Yao C, Wang Z, Zhao Z, Liu P, Yang F, Li X et al: Microbial characteristics across different tongue coating types in a healthy population. J Oral Microbiol 2021, 13(1):1946316.

9. Cui J, Cui H, Yang M, Du S, Li J, Li Y, Liu L, Zhang X, Li S: Tongue coating microbiome as a potential biomarker for gastritis including precancerous cascade. Protein Cell 2019, 10(7):496–509.

10. Jiang T, Guo XJ, Tu LP, Lu Z, Cui J, Ma XX, Hu XJ, Yao XH, Cui LT, Li YZ et al: Application of computer tongue image analysis technology in the diagnosis of NAFLD. Comput Biol Med 2021, 135:104622.

11. Hu J, Han S, Chen Y, Ji Z: Variations of Tongue Coating Microbiota in Patients with Gastric Cancer. Biomed Res Int 2015, 2015:173729.

12. Han S, Chen Y, Hu J, Ji Z: Tongue images and tongue coating microbiome in patients with colorectal cancer. Microb Pathog 2014, 77:1–6.

13. Lo LC, Cheng TL, Chen YJ, Natsagdorj S, Chiang JY: TCM tongue diagnosis index of early-stage breast cancer. Complement Ther Med 2015, 23(5):705–713.

14. Zhou G, Huang D, Cai Y, Huang K, Xie D: The Relationship Between the Characteristics of Tongue Manifestation and Clinical Classification in Patients with Coronavirus Disease 2019. Journal of Traditional Chinese Medicine 2020, 61(19):1657–1660.

15. Wang X, Wang X, Lou Y, Liu J, Huo S, Pang X, Wang W, Wu C, Chen Y, Chen Y et al: Constructing tongue coating recognition model using deep transfer learning to assist syndrome diagnosis and its potential in noninvasive ethnopharmacological evaluation. J Ethnopharmacol 2022, 285:114905.

16. Li J, Chen Q, Hu X, Yuan P, Cui L, Tu L, Cui J, Huang J, Jiang T, Ma X et al: Establishment of noninvasive diabetes risk prediction model based on tongue features and machine learning techniques. Int J Med Inform 2021, 149:104429.

17. Li J, Yuan P, Hu X, Huang J, Cui L, Cui J, Ma X, Jiang T, Yao X, Li J et al: A tongue features fusion approach to predicting prediabetes and diabetes with machine learning. J Biomed Inform 2021, 115:103693.

18. Li J, Huang J, Jiang T, Tu L, Cui L, Cui J, Ma X, Yao X, Shi Y, Wang S et al: A multi-step approach for tongue image classification in patients with diabetes. Comput Biol Med 2022, 149:105935.

19. Huo CM, Zheng H, Su HY, Sun ZL, Cai YJ, Xu YF: Tongue Shape Classification Integrating Image Preprocessing and Convolution Neural Network. 2017 2nd Asia-Pacific Conference on Intelligent Robot Systems (Acirs) 2017:42–46.

20. Hou J, Su HY, Yan B, Zheng H, Sun ZL, Cai XC: Classification of Tongue Color Based on CNN. 2017 Ieee 2nd International Conference on Big Data Analysis (Icbda) 2017:725–729.

21. Wang X, Liu J, Wu C, Liu J, Li Q, Chen Y, Wang X, Chen X, Pang X, Chang B et al: Artificial intelligence in tongue diagnosis: Using deep convolutional neural network for recognizing unhealthy tongue with tooth-mark. Comput Struct Biotechnol J 2020, 18:973–980.

22. Xu Q, Zeng Y, Tang W, Peng W, Xia T, Li Z, Teng F, Li W, Guo J: Multi-Task Joint Learning Model for Segmenting and Classifying Tongue Images Using a Deep Neural Network. IEEE J Biomed Health Inform 2020, 24(9):2481–2489.

23. Tang WJ, Gao Y, Liu L, Xia TW, He L, Zhang S, Guo JH, Li WH, Xu Q: An Automatic Recognition of Tooth-Marked Tongue Based on Tongue Region Detection and Tongue Landmark Detection via Deep Learning. Ieee Access 2020, 8:153470–153478.

24. Kanawong R, Obafemi-Ajayi T, Ma T, Xu D, Li S, Duan Y: Automated Tongue Feature Extraction for ZHENG Classification in Traditional Chinese Medicine. Evid Based Complement Alternat Med 2012, 2012:912852.

25. Zhou J, Li S, Wang X, Yang Z, Hou X, Lai W, Zhao S, Deng Q, Zhou W: Weakly Supervised Deep Learning for Tooth-Marked Tongue Recognition. Front Physiol 2022, 13:847267.

26. Li Z, Ren X, Xiao L, Qi J, Fu T, Li W: Research on Data Analysis Network of TCM Tongue Diagnosis Based on Deep Learning Technology. J Healthc Eng 2022, 2022:9372807.

27. Li S, Wang R, Zhang Y, Zhang X, Layon AJ, Li Y, Chen M: Symptom combinations associated with outcome and therapeutic effects in a cohort of cases with SARS. Am J Chin Med 2006, 34(6):937–947.

28. Miner AS, Milstein A, Schueller S, Hegde R, Mangurian C, Linos E: Smartphone-Based Conversational Agents and Responses to Questions About Mental Health, Interpersonal Violence, and Physical Health. JAMA Intern Med 2016, 176(5):619–625.

29. Yang Z, Zhao Y, Yu J, Mao X, Xu H, Huang L: An Intelligent Tongue Diagnosis System via Deep Learning on the Android Platform. Diagnostics (Basel) 2022, 12(10).

30. Hu MC, Lan KC, Fang WC, Huang YC, Ho TJ, Lin CP, Yeh MH, Raknim P, Lin YH, Cheng MH et al: Automated tongue diagnosis on the smartphone and its applications. Comput Methods Programs Biomed 2019, 174:51–64.

31. Hu MC, Cheng MH, Lan KC: Color Correction Parameter Estimation on the Smartphone and Its Application to Automatic Tongue Diagnosis. J Med Syst 2016, 40(1):18.

32. Jiang T, Hu XJ, Yao XH, Tu LP, Huang JB, Ma XX, Cui J, Wu QF, Xu JT: Tongue image quality assessment based on a deep convolutional neural network. BMC Med Inform Decis Mak 2021, 21(1):147.

33. Redmon J, Divvala S, Girshick R, Farhadi A: You Only Look Once: Unified, Real-Time Object Detection. Proc Cvpr Ieee 2016:779–788.

34. He K, Zhang X, Ren S, Sun J: Deep Residual Learning for Image Recognition. IEEE 2016.

35. Lu MY, Williamson DFK, Chen TY, Chen RJ, Barbieri M, Mahmood F: Data-efficient and weakly supervised computational pathology on whole-slide images. Nat Biomed Eng 2021, 5(6):555–570.

36. Zhuang Q, Gan S, Zhang L: Human-computer interaction based health diagnostics using ResNet34 for tongue image classification. Comput Methods Programs Biomed 2022, 226:107096.

